# Halves and fragments derived of tRNAs in Escherichia coli are selectively associated with 30S ribosomal subunits and in the cytosol

**DOI:** 10.64898/2025.12.02.691187

**Authors:** Eva Jacinto-Loeza, Augusto Uc-Mass, Gabriel Guarneros

## Abstract

The generation of tRNA halves and fragments (tsRNAs) has been associated with stressful growth conditions in eukaryotes, but reports on tsRNAs in bacteria are scarce. Here, we demonstrate that the presence of tsRNAs in Escherichia coli depends on active translation, and that they are found in the 30S ribosomal subunit and the ribosomal-free fraction, but not in the 50S subunits, 70S ribosomes or polysome fractions. However, upon dissociation into subunits at a low magnesium concentration, some of the tRNAs present in the monosomal and polysomal fractions are processed into tsRNAs. RNA-seq analyses tsRNA fractions revealed that all tRNA species in the cell were processed into fragment profiles that varied widely for each tRNA. The E. coli CP78 strain contains an unusually high concentration of tsRNAAsnGUU, but it is likely that only a fraction of this participates in translation. These tRNAs, along with others in the cell, were released from the ribosomes and processed into tsRNAs. The tRNAs in the ribosomal-free fraction appear to be cleaved by the same RNase that is active in ribosomes. We propose that tsRNAs are generated as an initial decay step for tRNAs remaining on ribosomes following translation arrest. However, tsRNAs may also have other functions.

## INTRODUCTION

The cleavage of tRNAs at the anticodon loop to produce tRNA halves and at other sites to generate tRNA fragments (tsRNAs) has been extensively documented in eukaryotic systems (1). Here, we refer to tRNA fragments of 30–35 nucleotides as ’halves’, and to those of approximately 20 nucleotides as ’tRFs’ (2). The canonical function of tRNAs in translation is to deliver specific amino acids to the ribosome for transfer to the growing polypeptide chain, as dictated by the succession of codons in the mRNA during translation. tsRNAs have been documented to have multiple biological functions, including acting as signalling molecules in the stress response, regulating gene expression in cell proliferation, suppressing translation, modulating the DNA damage response and contributing to neurodegeneration in eukaryotic systems (1). In these studies, tRNA processing has been associated with stressful conditions, such as oxidative stress in human, plant and Saccharomyces cerevisiae systems (5, 6), nutrient starvation in Tetrahymena thermophila and Giardia lamblia (7, 8), starvation during hyphae formation in Streptomyces coelicolor (9) and elevated pH in Haloferax volcanii (10).

In human B cells, a 22-nt-long fragment derived from tRNA^(Gly) and generated by the nuclease Dicer inhibits RPA1, an essential gene with the functional characteristics of a microRNA (11). In the archaeon Haloferax volcanii, a fragment derived from tRNAVal, formed following the alkaline treatment of cells, appears to target the small ribosomal subunit, thereby interfering with protein synthesis (10). Additionally, a 35-residue-long tRNAPro 5’ half, which is associated with ribosomes and polysomes, inhibits global translation in mammalian cells in vitro (12). In Escherichia coli, amino acid starvation is associated with the rapid degradation of cellular tRNAs; however, the production of tsRNAs has not been analyzed (13). The documented cases of tsRNAs generated in E. coli are associated with exogenous genetic entities; several of these are generated by cleavage at the anticodon loop. It has been demonstrated that Leptotrichia shahii CRISPR-Cas13a, when expressed in E. coli, elicits the cleavage of uridine-rich anticodon tRNAs in addition to targeting RNA degradation (14). Bacteriophage T4 infection activates the latent ribonuclease PrrC, which cleaves the wobble position of endogenous tRNALysUUU (15). Colicins D and E5 process specific tRNAs into halves by cleaving the anticodon loops (16). The MazF toxin from the toxin-antitoxin system of Mycobacterium tuberculosis exhibits endonuclease activity on the D-loop of tRNAPro14 and the anticodon loop of tRNALys43 (17). In Streptomyces coelicolor, tRNA cleavage has been observed in cultures grown on a minimal medium. All types of tRNA in this bacterium appear susceptible to cleavage, resulting in the accumulation of halves. However, cleavage is not significantly affected by amino acid starvation, stringent response induction or inhibition of ribosome function (9). The nucleases involved in tRNA cleavage have been identified as follows: Rny1 in eukaryotes (18); angiogenin in yeast (19); and Dicer and RNase Z in mammalian cells (20). In HeLa cells, Dicer is responsible for the processing of tRNAs, yielding abundant 5’-end fragments (21).

In this study, we analyzed the production of tRNA halves and tRFs in E. coli using Northern blot assays and RNA deep sequencing. We demonstrated that all types of tRNA in the cell undergo cleavage at the anticodon loop and, eventually, at other sites within the molecule. Some cleavage products remained associated with the 30S ribosomal subunit, suggesting that the processed tRNAs had participated in translation. Those that remained on paused ribosomes were released by ribosome disassembly and processed into halves and tRFs. Additionally, tRNAs in the ribosome-free fraction that may not have participated in translation were processed, with the resulting products being found in the soluble fraction of cell extracts. We speculate that anticodon loop cleavage may be the initial step in the tRNA decay pathway.

## MATERIALS AND METHODS

### Bacterial strains, growth conditions and cell lysis

The Escherichia coli CP78 strain (22; 23; Atherly and Menninger, 1972) was grown in one-litre batches of minimal medium (M9), supplemented with 0.4% glucose and 4 mg/100 ml of each of the 20 amino acids. Cultures were grown by shaking at 37 °C until the optical density at 600 nm reached 0.6. Then, 1 mM chloramphenicol was added for two minutes before the cultures were cooled on ice and spun down at 10,000 rpm for 10 minutes at 4 °C in a Beckman JA-14 centrifuge. The cell pellet was washed twice with 30 ml of pre-chilled buffer solution containing 10 mM magnesium chloride, 100 mM ammonium chloride, 20 mM Tris (pH 8.0) and 1 mM chloramphenicol, and then resuspended in 4 ml of pre-chilled buffer solution containing 0.1% NP-40, 0.4% Triton X-100, 10 U/ml RNase-free DNase I (Roche), 0.5 U/ml Superase In (Ambion) and 1 mM chloramphenicol. The suspension was quickly frozen in liquid nitrogen and cryogenically pulverized in a MM 400 Retsch mill using 50 ml jars and 25 mm grinding balls. The sample was pulverized in six cycles at 18 Hz for three minutes each, with the canisters being re-chilled in liquid nitrogen between cycles. The thawed lysate was spun down at 20,000 g for 10 minutes at 4 °C. The nucleic acid concentration in the clarified lysate was determined using a NANODROP 2000Spectrophotometer (Thermo Scientific) at 260 nm.

### Fractionation in sucrose gradients

Sucrose gradients ranging from 10 to 55% (w/v) in a buffer solution containing 1 mM of chloramphenicol (excluding the run-off experiment shown in Fig. 3) and 2 mM of dithiothreitol (DTT) were prepared using a Gradient Maker (Hoefer SG 50) in SW28 tubes (Beckman polyallomer centrifuge tubes, catalogue no. 344061). Samples were loaded onto the gradients and spun down in a Beckman SW28 ultracentrifuge rotor at 27,000 rpm for 5 hours at 4 °C. The gradients were then fractionated using an Auto Densi-Flow IIC (Buchler Instruments, Labconco Company). All gradient fractions were manually collected while continuously monitoring the absorption at 254 nm using an Econo-UV monitor (Bio-Rad).

### Treatment of fractions obtained from the gradient

Polysomes, 70S ribosomes, 30S and 50S ribosomal subunits, and ribosome-free fractions were collected from the sucrose gradients. These were then diluted to 19 ml with buffer and precipitated with cold absolute ethanol at a ratio of 1:1 (vol/vol). The ribosomal pellet was resuspended in 0.6 ml of pre-chilled buffer containing 0.5 U/ml Superase In and either 1 or 10 mM magnesium chloride, for disassembly or maintenance of translating ribosomes, respectively. The mixtures were incubated for 5 hours at 4 °C with gentle shaking and resolved on a second sucrose gradient to separate the components of the disassembled ribosomes. The final fraction products were treated as indicated above.

### RNA isolation under acidic conditions

Each ribosomal pellet was resuspended in 0.6 ml of a solution containing 10 mM sodium acetate (pH 4.5), 1 mM sodium ethylene-diamine-tetraacetic acid (Na-EDTA) and Superase In (Ambion). The RNA was extracted three times using equal volumes of hot acid phenol, acid phenol-chloroform and chloroform:isoamyl alcohol (24:1, v/v). The final aqueous layers were mixed with 2.5 volumes of ethanol and left on ice for 1–2 hours. The RNA was then recovered by centrifugation and dissolved in 10 mM sodium acetate (pH 4.5) and 1 mM Na-EDTA. The RNA concentration was determined by measuring the absorbance at 260 nm using a NANODROP 2000 Spectrophotometer (Thermo Scientific), prior to loading onto electrophoresis gels.

### For the Northern blot assay

Total RNA was resolved on 0.5 mm-thick, 4%/16% denaturing, bilayer polyacrylamide gels (PAGE) and transferred to Gene Screen Plus nylon membranes (PerkinElmer) using a Trans-Blot Semi-Dry Transfer Cell (Bio-Rad) at 400 mA for 40 minutes. The membranes were cross-linked using a UVC 500 cross-linker (Hoeffer). [³²P]-5′ end-labelled oligonucleotide probes against selected tRNAs were used to reveal all molecular species (full-length tRNAs and tsRNAs harboring 5′ ends) (see Supplementary Table 1). The membranes were hybridized overnight at 42 °C in a HB 1000 Hybridizer (UVP Laboratory Products). The sequences of all DNA oligonucleotide probes used in this study are summarized in Table 2.

### Deep sequencing of ribosome-free RNA fragments

We used 80–100 ng of 34–52 nt RNA fragments resolved through PAGE and excised from the gel (see Supplementary Fig. 1). Quantitation was performed using a Qubit™ RNA HS Assay Kit. We constructed a library essentially following the NEXTflex™ Small RNA-Seq Kit v3 (Bio Scientific) protocol. Inserts of 24 and 44 bp without adapters were eluted from acrylamide gels. The library was sequenced in two different configurations: first, in 38 cycles twice in a NextSeq 500 (Illumina), using a NextSeq 500/550 High Output Kit v2 (75 cycles) for fragments up to 35 nt long; and second, in 76 cycles once, using the same equipment kit, for fragments up to 76 nt long. The proportions of reads for the different types of RNA in the short and long fragments of the ribosome-free and 30S fractions were reproducible (see Supplementary Fig. 1).

### Bioinformatic processing

The raw RNA-seq files were analyzed using Fastqc v0.11.6 to check the quality of the reads (25). The sequencing read adapters were then removed using Cutadapt v1.9.1 (26), after which the reads were aligned to the Escherichia coli K-12 strain MG1655 genome (NC_000913.3) using Bowtie v1.1.2 (27). Bedtools v2.25 (28) was used to annotate and count the mapped reads. The SAM files obtained from alignment to the reference genome were processed with Samtools v1.19 to be visualized with GBrowse v2.54. Images related to the RNA-seq data were obtained with GBrowse. The sequences were filed in the NCBI database (https://www.ncbi.nlm.nih.gov/, accessed 29 August 2025; accession number PRJNA1312260).

## RESULTS

### tRNA halves and fragments were found in ribosome-free extracts, associated with 30S ribosomal subunits

During the analysis of small RNA species deep sequencing data from Escherichia coli CP78 (see below), we found that many RNA-seq reads closely matched the 5’ or 3’ end sequences of tRNA genes. To investigate the origin of these tRNA fragments, we searched for their presence in cell extract fractions. Cell extracts were separated using sucrose gradients to produce five fractions: ribosome-free, 30S and 50S ribosomal subunits, 70S ribosomes and polysomes. These fractions were analyzed using acid urea-PAGE and Northern blots with [32P]-labelled probes to detect full-length tRNA^(Arg)^(ACG), 5′ and 3′ tsRNAs (see Fig. 1 and Supp. Table 1 for deoxyoligonucleotide probe sequences). While all fractions showed variable amounts of full-length tRNA^(Arg)^(ACG), the tsRNAs present in the crude extracts (Fig. 1A) were only detected in the ribosome-free and 30S fractions (Fig. 1C and D). In some blots, a 5′ end fragment signal was detected in the 50S ribosome subunit fractions, which is probably due to leakage from the 30S fraction, as the tsRNA signal was only present in the 30S middle subfraction, which corresponds with the 30S OD peak (Fig. 1B, C and D, 30S.2). The 3′ and 5′ tRNAArgACG fragments were present in the ribosome-free and 30S fractions of the same samples. Note that the 3′ end signals of tRNA^(Arg)ACG in the ribosome-free samples migrated slightly above the 5′ end signals (see Fig. 1 and compare the ribosome-free lanes in panels C and D), indicating that the 5′ and 3′ halves are different sizes. Based on these results, we infer that at least some of the tRNA processing relates to ribosomal protein synthesis.

**Fig. 1.**
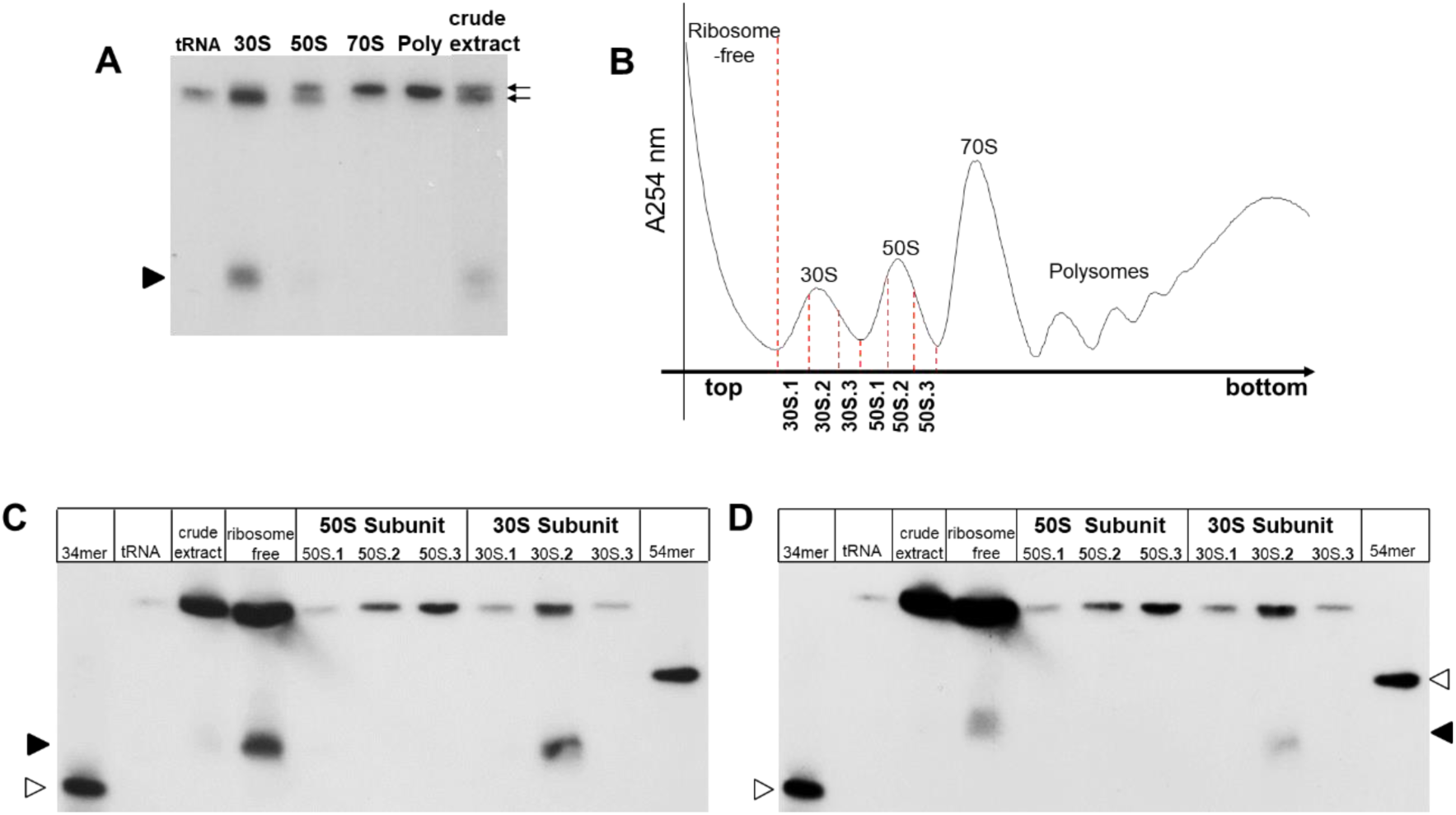
tRNA^Arg^ halves are present in the ribosome-free and 30S subunit fractions. **A)** Cell extracts of *E. coli* CP78 were resolved by a sucrose gradient, and the indicated fractions were analyzed by Northern blots with [^32^P]-5’tRNA^Arg^ACG probe. **B)** Sucrose gradient resolution of CP78 cell extract. **C** and **D)** Northern blots of the fractions revealing signals corresponding tRNA^Arg^ halves in the 30S and ribosome-free lanes. **C**) Northern blot revealed with the [^32^P]-5’tRNA^Arg^ACG probe. **D**) The nylon membrane used in **C**) was stripped-off 5’ probe and re-probed with [^32^P]-**3’**tRNA^Arg^ACG. The 30S and 50S fractions in **B** were split into three subfractions each (S.1 to S.3) to show that the faint signal in the 50S fraction in **A**) was leakage from the 30S fractions. The lane labeled ‘tRNA’ contained commercial pooled tRNAs. Full arrowheads indicate the positions of tRNA^Arg^ACG halves; open arrowheads indicate labeled oligonucleotide markers.

### A low magnesium concentration disassembles ribosomes and releases tRNA cleavage products

As shown above, 5′ and 3′ end tsRNAsArgACG were present in the 30S and ribosome-free fractions, but absent from the 50S, 70S and polysome fractions. However, full-length tRNA^(Arg)ACG signals were present in all fractions (see Fig. 1C and D). We suspected that the association of tsRNA with the 30S ribosomal fraction was due to the participation of tRNA in protein synthesis. To test this theory, the 70S and polysome fractions retrieved from a sucrose gradient (Figure 2A) were split in half. One half of each fraction was resuspended in a low magnesium concentration (1 mM) to disassemble the 70S and polysomes into 30S and 50S subunits (see Materials and Methods), while the other half was incubated in a high magnesium concentration (10 mM) to maintain the assembled translating ribosomes (31). The disassembled samples were separated using a second sucrose gradient (Figures 2B and 2C), and the resulting fractions were analyzed using a [32P]-5’tRNALeuCAG probe. Blots from the original gradient (Fig. 2D, lanes 3–5) showed at least two signals below the full-length tRNALeuCAG. These corresponded to the 5′ end of the tsRNALeuCAG in the ribosome-free and 30S fractions (Fig. 2, lanes 3 and 4). The signals detected in the 50S fraction (lane 5) were probably spilled over from the 30S fraction (see legend for Fig. 1). No detectable tRNA halves or tRFs were observed in the 70S fraction (lane 6). Interestingly, disassembly of the 70S and polysome fractions at a low magnesium concentration produced intense 5′ end tRNA^(Leu)CAG signals in the ribosome-free fractions (Figure 2D, lanes 7 and 11, respectively). No tsRNA signals appeared in the 30S or 50S fractions of the disassembled ribosomes (Figure 2D, lanes 8–9 and 12–13). These results suggest that the disassembly of ribosomes exposes tRNAs to an activity that cleaves them into halves and fragments. Furthermore, the generation of tsRNAs from tRNALeu is consistent with this processing being a general tRNA process.

**Fig. 2.**
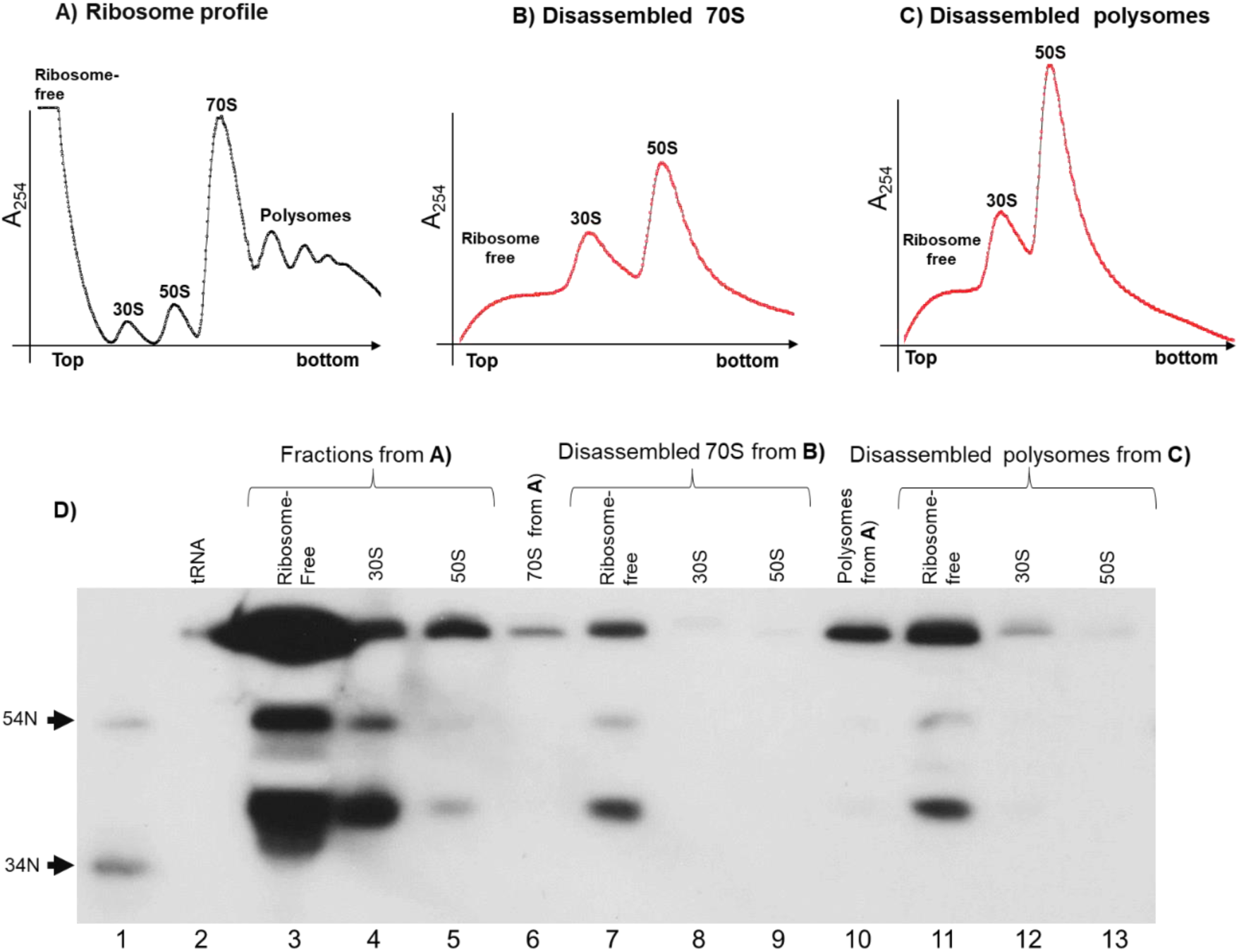
Dissociation of ribosomes paused in translation releases tRNA^Leu^ halves and fragments. **A**) One hundred OD260 units of crude extracts of *E. coli* CP78 grown on M9 medium were resolved through 10mM Mg^2+^ sucrose gradients. 70S **B**), and polysomes **C**), from **A**), were disassembled in 1mM Mg^2+^ and ran through second sucrose gradients. **D**) The fractions were resolved by denaturing PAGE and the tRNA fragments revealed by Northern blots with a probe specific for the [^32^P] 5’- tRNA^Leu^CAG. (See Material and methods for experimental details). The whole collected RNA fractions were loaded in each lane.

**Fig. 3.**
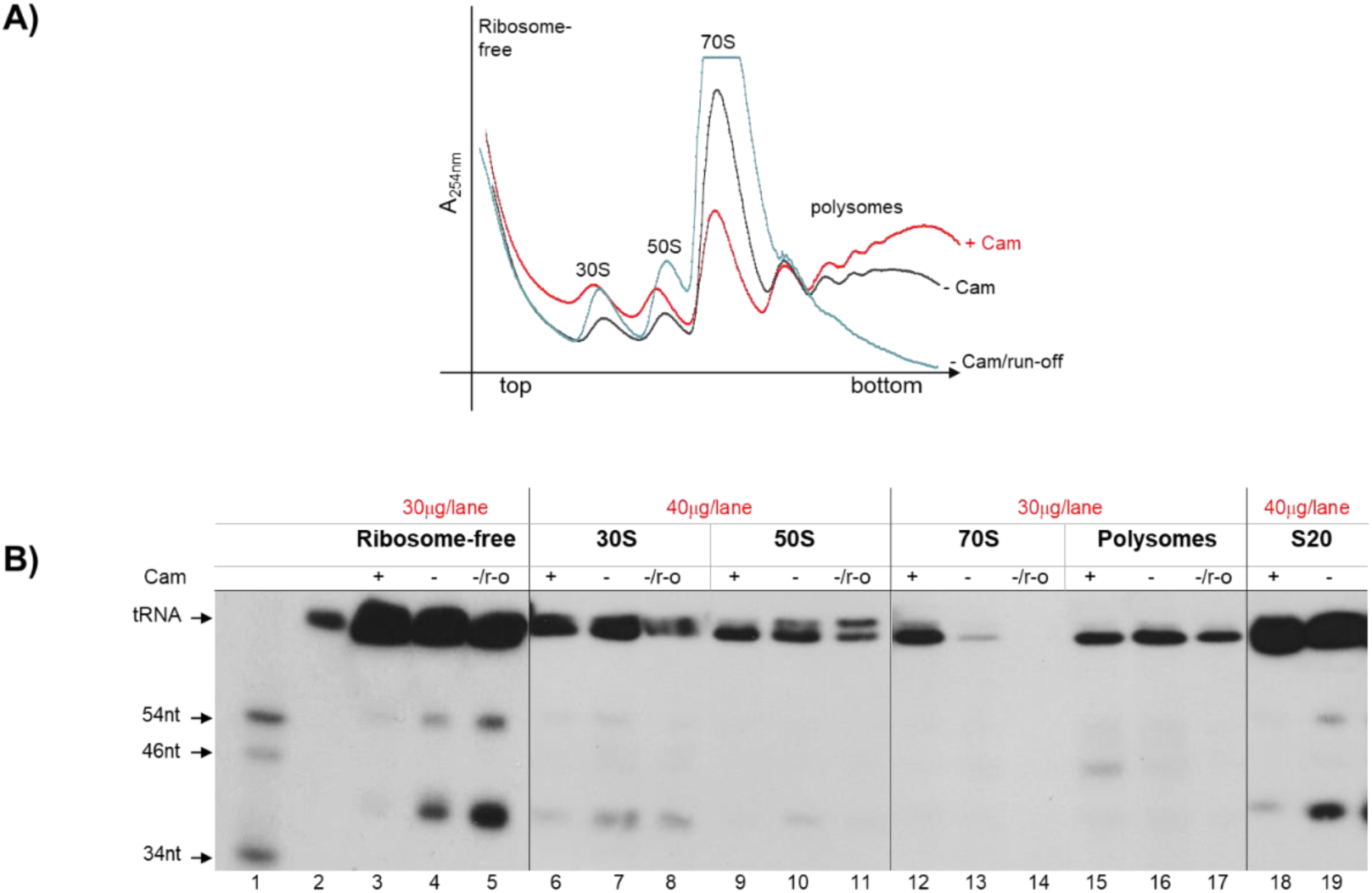
Production of tRNA fragments and halves correlates with active protein synthesis. **A)** Strain CP78 cultures were treated (+Cam) or untreated (-Cam) with chloramphenicol and cell-free extracts of each culture were prepared. An aliquot of the untreated cell extracts was supplemented with the appropriate reagents to allow in vitro run-off translation for 1h at 25°C (-Cam/run-off). The cell extracts of the three samples were resolved through sucrose gradients and fractions were separated. **B)** Northern blots of RNA isolated from extracts treated with chloramphenicol (+), untreated (-) and untreated plus run-off (-/r-o). The RNA from the collected fractions in **A** was ran though acid urea-PAGE, blotted and hybridized with a [^32^P]-5’ tRNA^Leu^CAG probe. Labeled deoxi-oligo nucleotides were included as size markers in lane 1; lane 2 contains a commercial preparation of *E. coli* tRNA. Lanes were loaded with the indicated amounts of RNA. Film time exposures of the different panels were adjusted to visualize halves and fragments.

### The production of tRNA fragments and halves is dependent on active protein synthesis

Our previous results suggest that tRNAs remaining on ribosomes that have become arrested during translation are protected from cleavage. However, cleavage occurs when the tRNAs are released upon dissociation of the ribosomes into subunits. We investigated whether inhibiting protein synthesis with chloramphenicol affects tRNA processing. Chloramphenicol (Cam) inhibits translation by binding to the A-site of the peptidyl transferase center in the ribosome, thereby preventing aminoacyl-tRNA from accepting the nascent peptide during translation elongation (32, 33, 34). Chloramphenicol was added to bacterial cultures for two minutes and then ice-chilled to arrest protein synthesis before the cells were harvested. Then, it was added at different stages during cell extract preparation (see Materials and Methods). We monitored the tsRNALeuCAG sequence signal to illustrate what may have happened to other tsRNAs. Cam treatment reduced the accumulation of 70S ribosomes and increased polysome formation (Fig. 3A), as well as decreasing tsRNA signals in the ribosome-free and 30S fractions relative to untreated cells (Fig. 3B; compare lanes 3 and 4 and lanes 6 and 7). We concluded that tRNA processing depends on active protein synthesis. These results suggest that tRNA cleavage occurs when tRNAs are released from translating ribosomes. To test this hypothesis, we divided the Cam-untreated cell extracts into two portions: one was supplemented with reagents for run-off translation, while the other was kept as a control. It was expected that the run-off translation would proceed with the termination and release of tRNAs and mRNAs (see Materials and Methods). Run-off translation reduced the polysome concentration and increased the accumulation of 30S, 50S and 70S ribosomal particles (see Fig. 3A). However, the signals for tsRNAs were more abundant in the ribosome-free fractions of run-off-treated samples than in untreated samples (Fig. 3B; compare lanes 5 and 4). This final result suggests that, once translation is terminated and ribosomes are released from mRNA, tRNA cleavage also occurs.

In addition to chloramphenicol treatment, we evaluated the effect of temperature upshifting on translation in an E. coli pth(ts) mutant (23); (36); Suppl. Fig. 4). Probing for tRNALeuCAG and tRNAAlaUGC sequences showed that tsRNA production gradually decreased with incubation time at a non-permissive temperature in a pth(ts) culture. These results are consistent with the observation that tsRNA production depends on protein synthesis for all tRNAs.

### All tRNA species in the cell are present in the ribosome-free and 30S fractions in the form of halves and fragments

We have demonstrated the existence of tsRNAs that are homologous to both tRNAArgACG and tRNALeuCAG within the ribosome-free fractions, as well as within the 30S-associated fractions, of cell extracts (see Figures 1 and 2). To investigate whether these fractions contained tsRNAs from other tRNAs, we conducted a deep RNA-seq experiment (see Materials and Methods). The RNAs were separated by acid urea PAGE and the gel areas containing small RNAs between 34 and 54 nucleotides in length were excised. The RNA was then purified and treated for RT-PCR deep sequencing (see Materials and Methods). The presence of different types of cellular RNA, including ribosomal, messenger and non-coding RNA, was detected in both the ribosome-free and 30S fractions (see Supplementary Fig. 2A and B). Data for the overall RNA-seq reads versus nucleotide lengths were plotted for the different types of RNA (see Supplementary Fig. 2C). A 36 nt column of tsRNA reads stands out in the ribosome-free fraction, which is absent from the 30S fraction.

This excess of tsRNA reads derives from tRNAAsn, as it is radically reduced when reads homologous to tRNAAsn are filtered out informatively (not shown). Conversely, the percentage of tsRNA sequences was lower in the 30S fraction than in the ribosome-free fraction (see Supplementary Figures 2A and 2B), as expected given the high relative contribution of ribosomal RNA in the 30S fraction. However, the actual levels of specific tsRNAs derived from tRNAArg (Fig. 1) and tRNALeu (Figs. 2 and 3), as analysed by Northern blots, were also higher in the ribosome-free fraction than in the 30S fraction.

In the RNA-seq experiment, the number of tsRNA reads in the ribosome-free fraction was higher than in the 30S subunit fraction, even when the tRFs and tRNA halves homologous to tRNAAsn were subtracted (see Supplementary Table 2 for total data, columns 2 and 3). Plots of size against the number of tsRNA reads obtained for both fractions suggest that two tsRNA populations exist, peaking at 29 nt (tRFs) and 35 nt (tRNA halves) (Fig. 4 A and B). Since the fractions contain comparable tsRNA length distributions, albeit at different proportions, it is possible that the tRNA processing activity would be identical, but with different rates of further decay for both fractions. The ratio of tRNA halves to tRFs is about 13-fold higher in the ribosome-free fraction than in the 30S fraction (Fig. 4). This disparity probably reflects differences in the tRNA decay regime for each tRNA in the fractions. The proportion of 5′ to 3′ tRNA halves was 3:1 in the ribosome-free fraction, compared with a 1:1 ratio in the 30S fraction (see Supplementary Figures 3A and 3B). We speculate that this difference is due to the tRNAs being protected against nuclease activity while they remain associated with ribosomes. As has been proposed for stable RNAs, it is likely that tRNA degradation is initiated by endonuclease cleavage, followed by exoribonuclease in a stepwise manner (37).

**Fig. 4.**
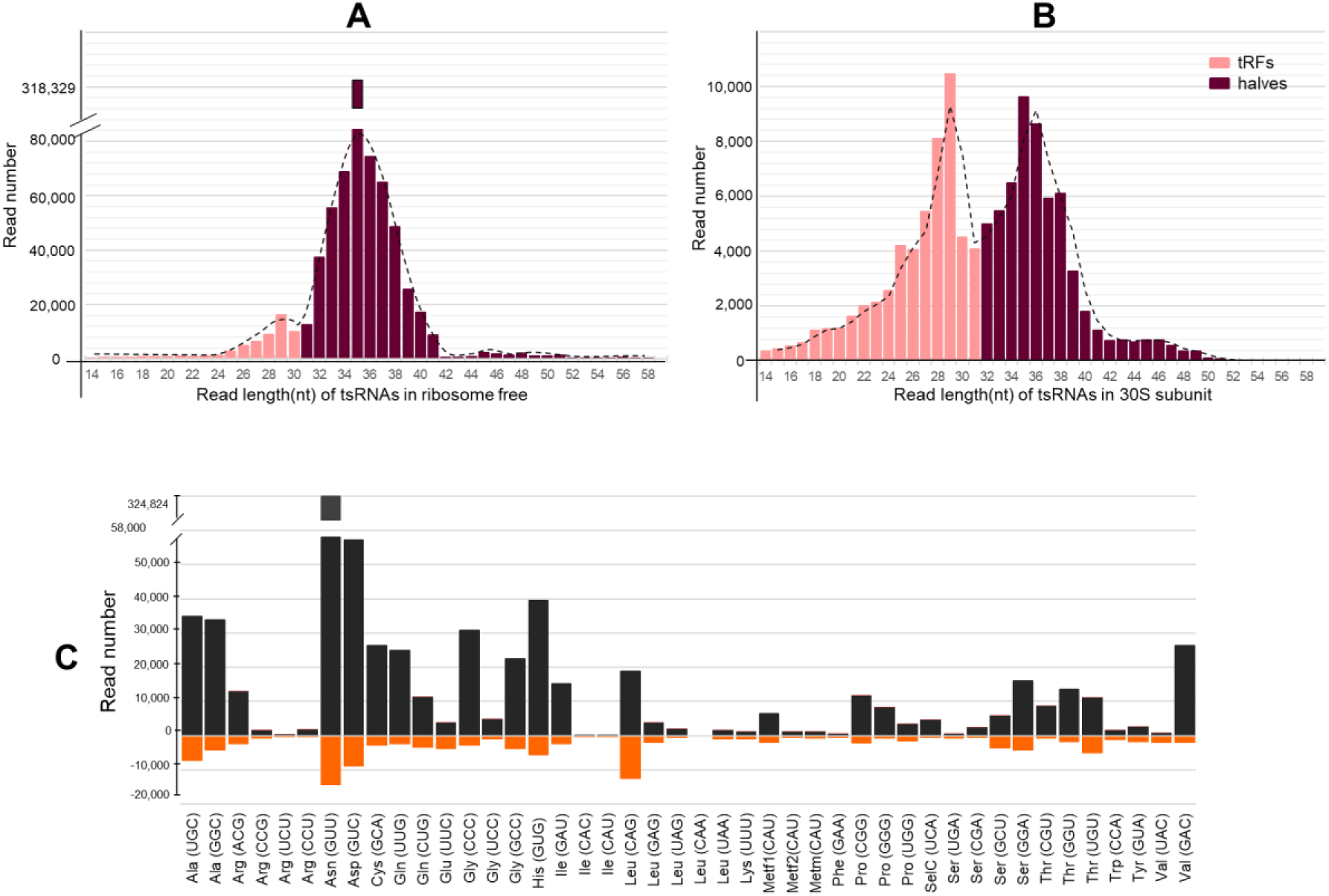
Proportion of RNA types and size distribution of tRFs and tRNA halves in the ribosome-free and 30S fractions. Length distribution of tRNA seq reads by nucleotide numbers of data sets obtained for **A)** ribosome-free and **B)** for 30S subunit tRNA fractions. The plots show tRNA-derived sequences cresting at 29nt (tRFs) and 35nt (halves). **C)** Quantity correlation between RNA seq reads of tRFs and halves in the ribosome-free and 30S fractions. tRNA-seq reads are plotted on mirror scales for comparison; ribosome-free data (black bars in the positive scale) correspond to reads in 30S fraction (orange bars in the negative scale). The reads were assigned to each tRNA by sequence homology with tRNAs of *E. coli* (NC_000913.3). Note that tRNA^Asn^ reads for the ribosome-free fraction are about 7-fold more abundant than those for the 30S fraction (Supp. Fig. 1C). The corresponding tRNAs in the horizontal axis are identified by anticodon.

### Correlation between tRNA fragments in the ribosome-free and 30S fractions, and an excess of tRNAAsnGUU in strain CP78

The number of RNA-seq reads for each tRNA in the ribosome-free and S30 subunit fractions were compared (see Fig. 4C). Most tsRNA reads in the ribosome-free fraction correlate directly with the 30S subunit fraction reads, except for tsRNAAsnGUU, which shows an extremely high number of reads in the ribosome-free fraction. This correlation can be easily visualised by overlaying the tendency plots for the normalised number of reads for the 5’ and 3’ halves of each fraction. (Supplemental Figures 3C and 3D). It is unlikely that all of the excess tRNAAsnGUU participates in translation. Typically, tRNA abundance is proportional to codon usage, but asparagine codons are not particularly abundant in E. coli (38). We suggest that some tRNAAsnGUU involved in protein synthesis remained attached to ribosomes, was released into the solution, and was cleaved into tsRNAs when the ribosomes dissociated into subunits. However, it seems that a high proportion of the tRNAAsnGUU pool is processed into tsRNAs. We offer two alternative explanations for this observation: either tsRNAs in the 30S fraction are released and accumulate in the ribosome-free fraction, or a unique ribonuclease is responsible for processing tRNAs separately in both the ribosome-free and 30S fractions, or some other complex degradation process occurs.

### The cleavage pattern may depend on the nucleotide composition of tRNAs and on whether they are associated with the ribosome

The GBrowse viewer was used to generate a collection of tRNA sequence reads that are homologous to each of the 81 tRNA genes that were identified in Escherichia coli CP78 (RefSeq NC_000913.3). Cleavage profiles and read lengths for each tRNA were obtained for the 30S and ribosome-free fractions (Fig. 1; Supp. Table 2), which contain tRFs and halves. The tsRNA profiles for each tRNA differ, but most show a gap of variable length missing nucleotides in the anticodon loop region. The profiles also demonstrate uneven abundance of 5′ and 3′ end halves (see Fig. 5 and Supp. Table 3). Some profiles show an abundance of 5′ end halves in both the 30S and ribosome-free fractions, such as for most of the tRNASerGCU (serV gene) and tRNALeuCAG (leuT gene) profiles. Others, such as the tRNAArgCCU (argW gene) profile, show a greater number of reads for the 3′ end halves in both fractions. Some tRNAs appear to undergo processing, generating a central fragment that produces a pyramidal profile for tsRNAs of tRNAMetf2CAU (metY gene) and tRNAValUAC (valZ gene) in the ribosome-free fractions, as well as partial gap cleavage in the 30S fraction (see Fig. 5). These results may reflect different cleavage patterns, as well as the partial protection of tRNAs through their association with ribosomes, and the secondary ribonuclease decay of the 5′ and 3′ halves, depending on the structure of the tRNA and its nucleotide modifications. In all tsRNA sequences visualised by GBrowse, a nucleotide is missing at the cleavage site. We assume that the missing nucleotide in the tRNA sequence profiles corresponds to cleavage generating tsRNAs. However, we cannot determine whether cleavage occurred at the 5′ or 3′ bonds of the missing nucleotide, or at both. We do not know whether the missing nucleotide is an artefact of the GBrowse viewer or whether it has been lost due to nuclease activity. For example, the tRNA^(Glu)UUC (gltT gene) the sequence often exhibits a one-nucleotide gap at the anticodon UUC (the missing nucleotide is indicated by brackets) or at variable sites within the anticodon loop (see tRNA-Glu-UUC, GltW in NCBI accession number PRJNA1312260).

**Fig. 5.**
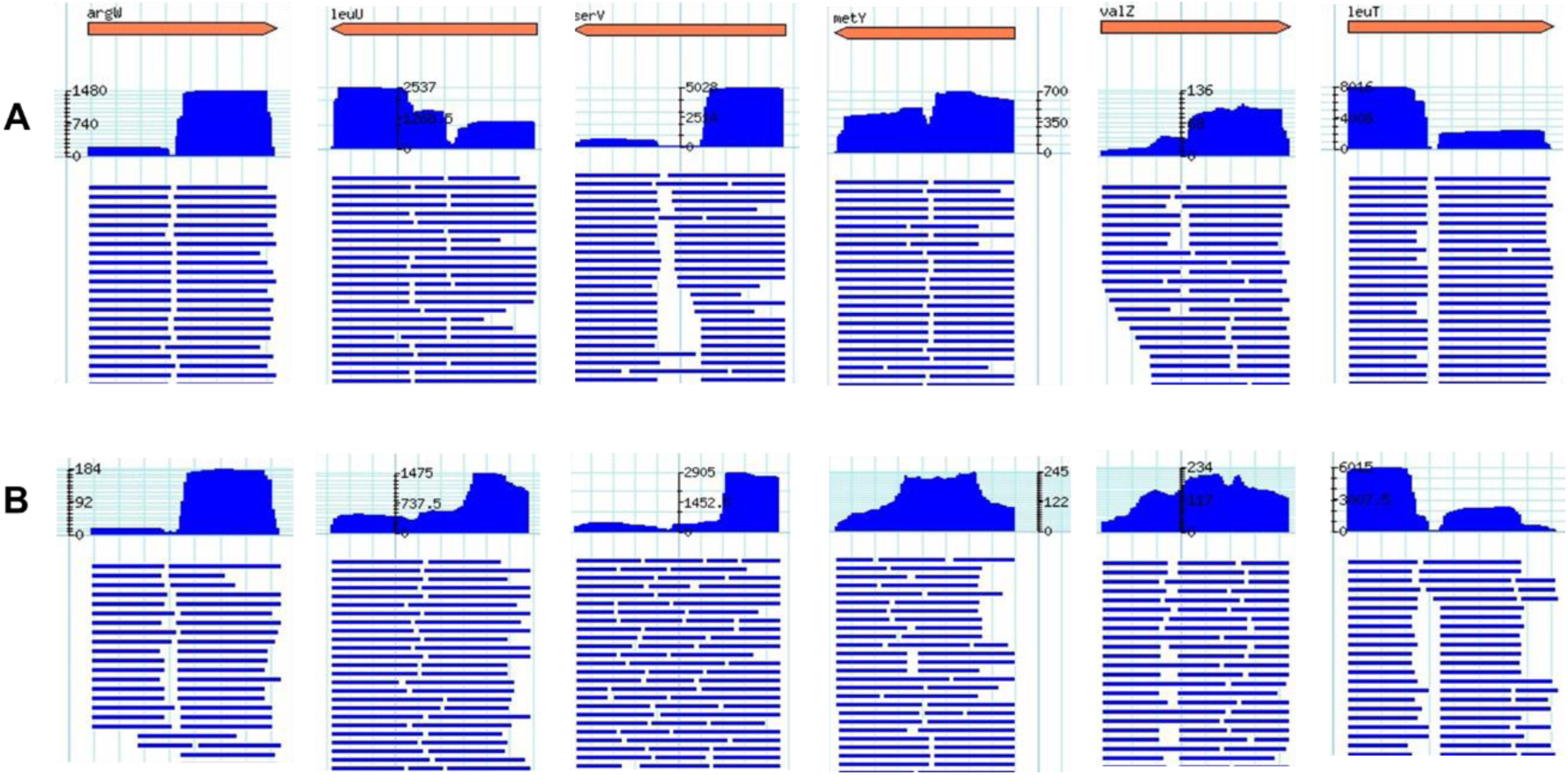
Coverage of RNA reads for halves and fragments of selected tRNAs. **A)** for the ribosome-free fraction, **B)** for the 30S fraction. The arrows in orange at the top represent the tRNA genes; arrowheads indicate 3’ ends. The arrows conserve the orientation of the genes in the bacterial chromosome. The histograms (blue) in each panel represent abundance and size of the corresponding RNA reads according to the embedded scales and gene orientations. The gaps that split most profiles in about halves indicate the site of cleavage at the anticodon loop. Other cleavage sites generate multiple fragments. Note that some tRNA reads show mainly abundance either for 5’ or 3’ halves in both ribosome-free and 30S fractions. Images generated by G-Browse.

In some cases, cleavage has occurred at a secondary site, resulting in the creation of an internal tRNA fragment, as seen with the tRNALeuGAG leuU gene (Fig. 5A). This tRNA contains a long variable loop, and a second cleavage could have occurred, as the predicted fragments corresponded to the expected fragment size. However, the secondary structure of tRNALeuCAG (leuT gene) also features a long variable loop, yet it does not appear to undergo frequent cleavage (Fig. 5). We do not yet know what elements affect cleavage at tRNA loops. Nevertheless, another cleavage seems to occur proximal to the 3’-end of the leuT gene tRNA, probably at the ∝ loop (Fig. 5B). Missing nucleotides were recognised at different locations, and within the anticodon loop they may be inherent to site cleavages or created by exonucleolytic decay.

## DISCUSSION

In this study, we present evidence of the formation of tsRNAs (tRNA halves and tRFs) and their presence in the 30S and ribosome-free fractions of cellular extracts. tsRNA generation occurs for all tRNA species present during active translation, as demonstrated by deep RNA sequencing of fragments. The dissociation of ribosomes that are arrested during translation or translation termination is accompanied by the release of tRNAs and their specific cleavage, which suggests that these tRNAs are substrates for decay (13). Low magnesium concentrations appear to disassemble ribosomes paused in translation and activate an RNase associated with the ribosomes. However, our evidence also suggests that tRNAs in the cytosol are processed into tsRNAs. Cleavage of excess tRNAAsn in the ribosome-free fraction that presumably did not participate in ribosomal translation also appears to be a substrate of the processing activity. This cleavage activity may be RNase I, as it has been reported that this activity is inhibited by magnesium and is associated with 30S ribosomes (39). The processing observed here may represent the initial steps in a decay pathway for tRNAs that are unable to participate in further translation cycles, or the generation of tRNA halves that may serve as a source of secretion biomolecules (40). The different conditions used to arrest translation reduced and eventually stopped the generation of tsRNAs. In addition to chloramphenicol and ice treatments, translation was arrested by shifting a growing culture of CP78 pth(ts), a thermosensitive variant of peptidyl-tRNA hydrolase, to a non-permissive temperature (Suppl. Fig. 4; (23); (36).

Deep sequencing experiments were conducted on small RNA fragments isolated from ribosome-free and 30S fractions (see Supplementary Figure 1). These experiments showed that the tsRNAs matched all the tRNAs present in strain CP78. Of the 43 tRNAs that were positively identified by sequence, 42 showed tsRNAs (see Supplementary Table 2). Sequences corresponding to tsRNALeu4 were absent, probably due to a gene deletion, as previously described (41). In vivo, most tRNAs on ribosomes are in the post-state, i.e. after the tRNA has been translocated from the A and P sites to the P and E sites. The tRNA in the E site is deacylated and ready to be released from the ribosome, whereas the tRNAs in the P site are peptidylated (42). The observation of two distinct tRNA bands in Northern blots of the 50S subunit from Cam uninhibited translation (Fig. 3, lanes 10 and 11) suggests that they may correspond to free aminoacylated or peptidylated tRNA. The presence of chloramphenicol, which was used for most of the RNA extractions, inhibited protein synthesis, thereby preventing the formation of peptide bonds. Alternatively, in aborted 50S subunits, the translocation of the tRNA moiety from the A site to the P site frees up the A site, allowing a release factor to cleave off the tRNA (43). In the present case, it is also possible that the A site could be occupied by an aminoacyl-tRNA.

The accumulation of tsRNAs has been attributed to different types of stress in other organisms. The data presented here suggest that the ultimate cause of tsRNA generation in these cases may be the processing of tRNA upon translation arrest. For example, protein synthesis was eventually arrested due to nutrient starvation in Tetrahymena thermophila (7), Giardia lamblia (44) and Streptomyces coelicolor (9). The increased concentration of a tRNAAspGUC fragment under methionine starvation in an S. cerevisiae methionine auxotroph may also result from the inhibition of protein synthesis (5). Other stress treatments not directly related to the inhibition of protein synthesis, such as H₂O₂-induced oxidative stress in Saccharomyces cerevisiae, result in the inhibition of translation by interfering with initiation, elongation, and/or termination. However, stress conditions such as UV irradiation or starvation for glucose or nitrogen in yeast did not increase tsRNA accumulation (5). Further research is needed to clarify this issue.

The association of tsRNAs with the 30S ribosomal subunit suggests that processed tRNAs are involved in ribosome translation and remain unaffected by endonuclease activity as long as they are coupled with ribosomes (see Fig. 2, lanes 6 and 10). The association of tsRNAs with the 30S subunit, but not the 50S subunit, may be related to the affinity of tsRNAs for the 30S subunit (46). It is possible that some of the tsRNAs that accumulate in the ribosome-free fraction originate from tRNAs that are engaged in translation and released and processed by ribosome dissociation after translation termination (Fig. 1C and D). During run-off translation, the concentration of stalled ribosomes harbouring tRNAs is low, presumably because most terminated translations release the last-used tRNA (47, 48). A run-off experiment, in which protein synthesis continues until one of the reagents is depleted, revealed an increase in tsRNALeuCAG tsRNAs in the ribosome-free fraction, but not in the 30S fraction (Fig. 3). This result suggests that ribosomal translation generates tsRNAs that are released into the cytosol. This implies that tsRNAs in the soluble fraction may originate from ribosome drop-off and direct processing of free tRNAs. Although some tRNALeuCAG remained on 70S monosomes and polysomes, no halves or fragments were associated with them (Fig. 3A, lanes 6 and 10). However, disassembly of ribosomes at a low Mg²⁺ concentration released some of the tRNA, which was cleaved into halves and fragments (Fig. 2, lanes 7 and 11). This observation suggests that tRNALeuCAG and probably other tRNAs, once free in solution, are cleaved by a nuclease activity associated with the ribosome fraction. In other words, the tRNAs coupled to the ribosomes are inaccessible to the processing activity.

The dissociation of ribosomes and polysomes induced by low magnesium concentrations is accompanied by the release of tsRNAs, presumably as processing products of tRNAs that remain on ribosomes that are arrested in translation (see Fig. 2D, lanes 7 and 11). Regarding the tsRNAs in the ribosome-free fractions (Fig. 2D, lane 3), two non-exclusive explanations are considered: i) they originate from tRNAs that are cleaved on the ribosomes and released into the cytosol, as mentioned above, or ii) the tRNAs in the cytosol undergo direct processing by an endonuclease activity that is identical to that present on the ribosomes. Indirect support for the latter possibility comes from the unusually high level of tRNAAsnGUU in strain CP78, which is much higher than would be expected based on codon usage (see Figs. 4 and 5; references (38) and (49)). Only a fraction of the excess tRNAAsnGUU is expected to take part in translation; therefore, most of it may remain in the cytosol and be processed into tsRNAs by an activity identical to that present on the ribosomes. Indeed, most tsRNAAsnGUU was found in the ribosome-free fraction (see Figs. 4A and 4C). An unusually high number of tsRNA reads have been reported for tRNAGlu and tRNAAsp in E. coli (40), as well as for tRNAValGAC in the archaeon Haloferax volcanii. In the latter case, a 26-nucleotide-long tRF from the 5′ end of tRNA^(Val)(GAC) was by far the most abundant fragment quantified, implying that tRNA^(Val)(GAC) is hyperabundant in this strain (10).

Although the proportions of tRNA halves and 5’ to 3’ halves associated with the 30S fraction are similar, this is not the case for the ribosome-free fraction (see Supplementary Figures 3A and 3B). The proportions vary widely for each individual tRNA (see Supplementary Table 2). The more abundant tRNA half could be either the 5’ or 3’ end, depending on the specific tRNA and whether the analysed fraction was ribosome-free or 30S (see Supplementary Table 3). It is possible that the relative abundance of tRNA halves and tRFs is the result of initial anticodon loop cleavage, followed by secondary decay via other endo- or exo-RNase activities, as well as the structural accessibility of the RNAs to degradation (10).

We lack direct evidence regarding the nature of the nuclease activity that processes the tRNAs into tsRNAs. However, several observations made here are compatible with those made on RNase I by other authors. Although most RNase I is found in the bacterial periplasmic space, it is also associated with the 30S subunit of the ribosome, where its activity is inhibited by helix 41 of the 16S RNA (50). This is consistent with our observation that tRFs and tRNA halves are associated with 30S subunits (Fig. 2). Low magnesium concentrations disassemble ribosomes into subunits and also activate RNase I. In addition, the generated 3′ halves lack two terminal nucleotides of the amino acid acceptor sequence CCA (51). Interestingly, some of the 3′ tsRNAs in the sequencing data visualised here by GBrowse lack one or more of the 3′ CCA end nucleotides in halves produced from cleavage at the anticodon loop. It is unclear whether the anticodon cleavage and the absence of terminal 3’ CCA nucleotides are brought about by the same nuclease activity.

## Acknowledgements

We would like to thank Rogelio Cruz-Vera for his careful review of an early draft of the manuscript and Ricardo Grande for conducting the deep RNA-seq experiments included here. We would also like to thank Jair Martínez, Flor Revillas and Gaby Mora for their technical support. This work was supported by Secihti FC 2026/1602 grant, Mexico (formerly Conahcyt).

