## Supplementary Information for "Halves and fragments derived of tRNAs in Escherichia coli are selectively associated with 30S ribosomal subunits and in the cytosol"

V. Nov. 2025

Departamento de Genética y Biología Molecular, Centro de Investigación y de Estudios Avanzados del IPN. Apartado Postal 14-740, México

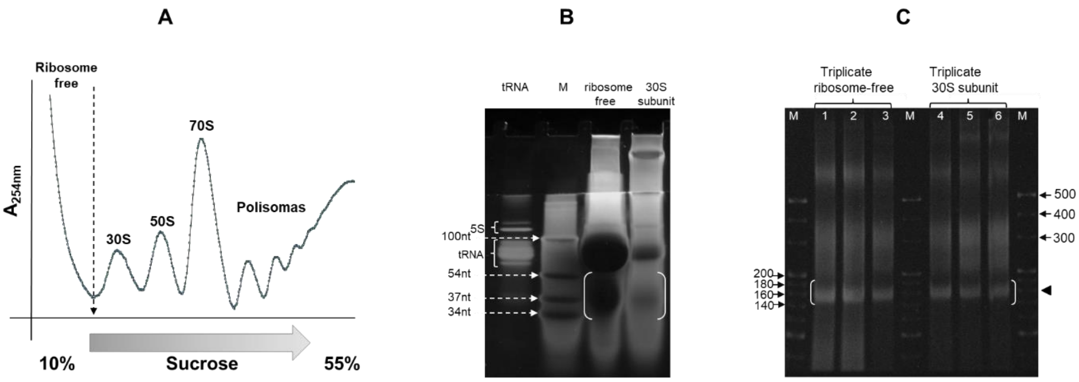


**Suppl. Fig. 1 Isolation of tRNA fragments and halves for RNAseq. A**) Ribosome profile of a sucrose gradient of cell lysates prepared from exponentially grown *E. coli* CP78. **B**) RNA fragments were resolved by acid urea PAGE from the 30S ribosomal subunits and ribosome-free RNA fractions and stained with SYBR Gold. The region in brackets was excised from the gel and treated for RNAseq. A mix of deoxy-oligos of known size was used as markers (lane M, white arrows). **C**) Triplicates of RT-PCR-amplified libraries for sequencing were resolved by nondenaturing 8% PAGE and loaded with the indicated PCR reactions (lanes 1 to 6). The brackets indicate the cDNA of tsRNA used for sequencing. M lanes are commercial DNA markers of the indicated nucleotide sizes.

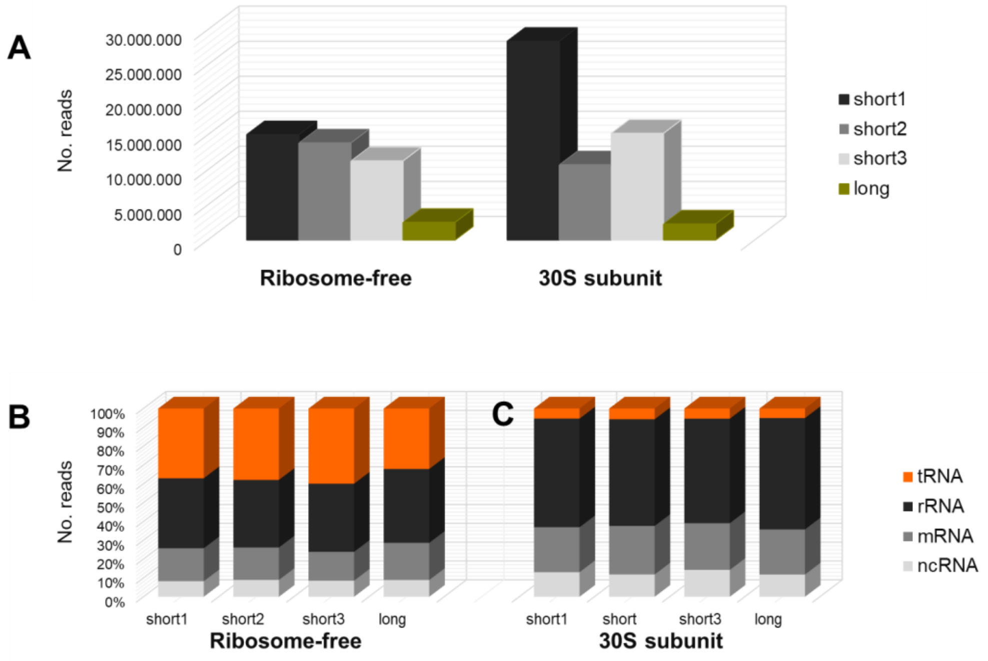


**Suppl. Fig. 1'. Reproducibility of the proportions of RNA types in the ribosome-free and 30S cell fractions**. Two RNA-seq experiments were done: one by triplicate selecting for short reads (14 to 35 nt), and another one selecting for long reads (14 to 57 nt). The indicated fractions were excised from RNA electrophoresed samples (see Suppl. Fig. 1, B and C). **A)** Histograms showing the overall number of reads obtained in the experiments that were used to estimate the indicated percents of each RNA type in the ribosome free fractions (RF) **B)** and in the 30S fractions **C)**.

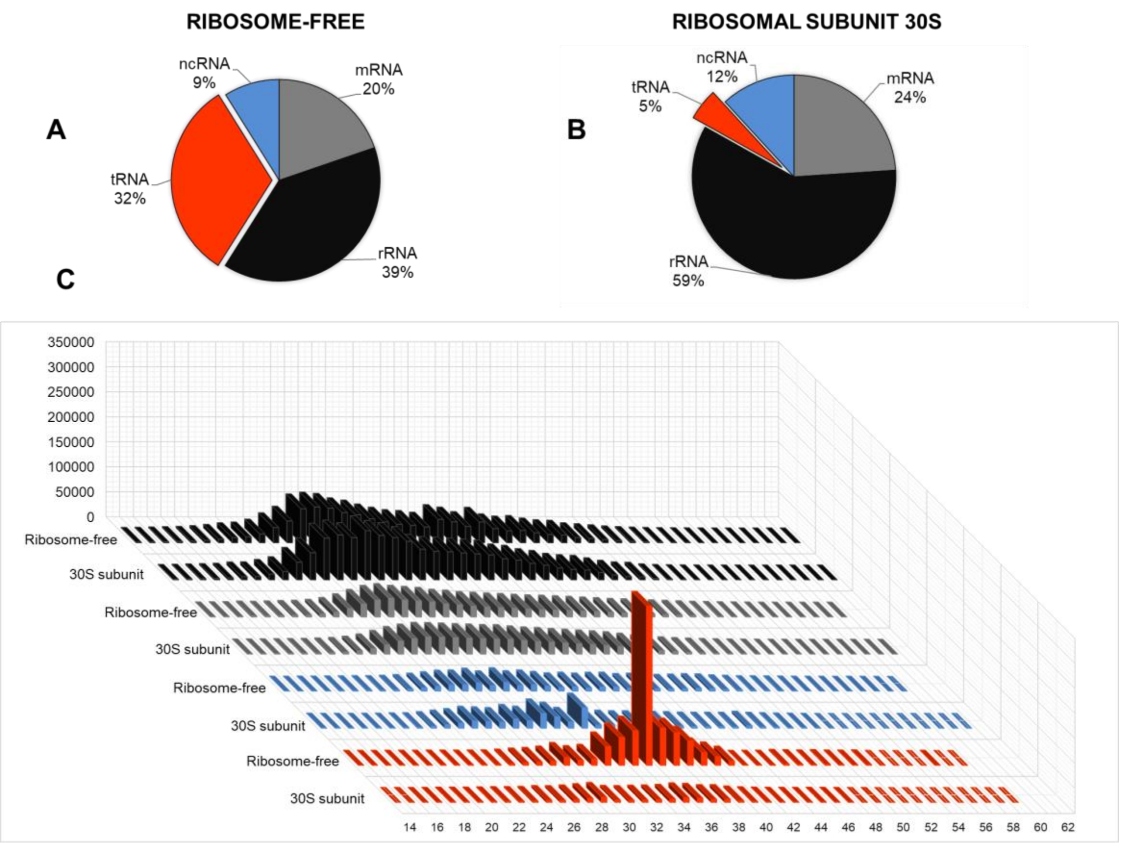


**Suppl. Fig. 2 Fractions and size distribution of tRFs and tRNA halves (tsRNAs) in the ribosome-free and 30S fractions.)** Pie charts showing the percent of RNAseq reads mapping to the different RNA types in data sets obtained for the ribosome-free **A),** and 30S RNA **B),** sucrose gradient fractions, respectively. **C**) Length distribution of reads in the ribosome-free, and 30S fractions. The types of RNA in **A**, **B** and **C** are color coded. The fractions were obtained from strain CP78 cell-free extracts and resolved through a sucrose gradient (see Suppl. Fig. 1). Data are from the long fragments sequencing (76 nt long, see Suppl. Fig. 5). Shown here are the reads that matched the map to the *E. coli* K-12 MG1655, NC_000913.3 genome annotated as tRNA, mRNA, rRNA and non-coding RNA (ncRNA). The tallest reads column in the ribosome-free tRNA histogram corresponds to the excess tRNA$^{Asn}$GUU (see text).

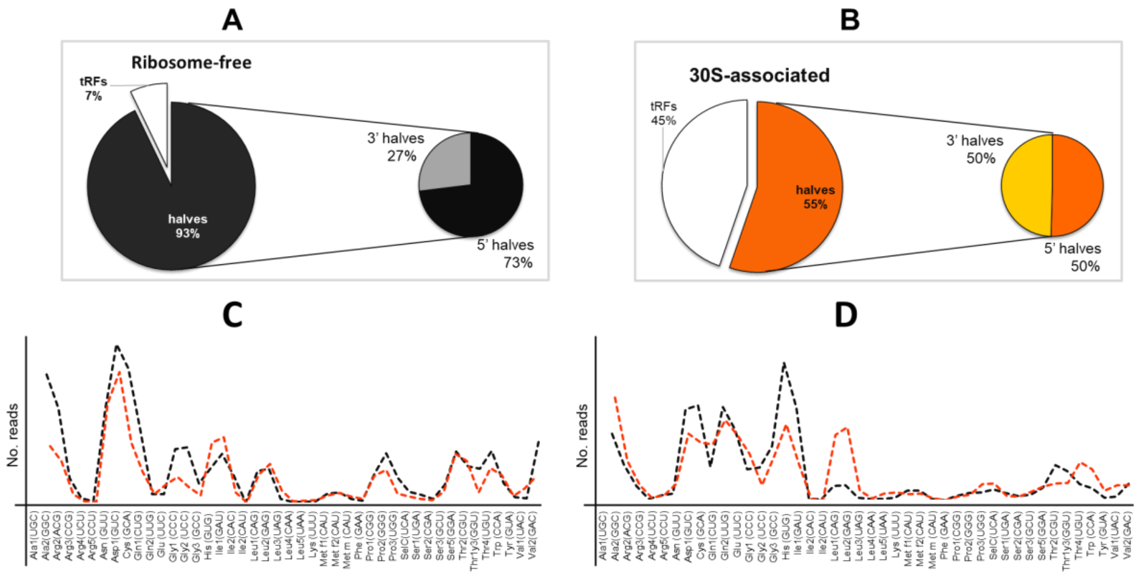


**Suppl. Fig. 3 Distribution and frequency of tsRNAs reads in the ribosome-free and 30S fractions.** Percent of RNAseq reads for overall 5' and 3' tRNA halves and tRFs in **A**) for the ribosome-free fraction, **B**) 30S ribosomal subunits fraction. **C**) Read tendencies overlap for 5'-tRNA halves. **D**) Tendencies overlap for 3'-tRNA halves. Tendencies for the ribosome-free fraction (black) and for the 30S fraction (orange) were plotted using Microsoft Excel (2016), with reads adjusted for overlapping. The tsRNAs sequencies largely matched tRNAs sequences ordered alphabetically in the horizontal axis. tRNAs are identified by amino acid specificity, isoacceptor number and anticodón sequence. The relative number of reads were based on those in Supp. Table 2.

| Name | Target | Purpose | Sequence (5'-3’) | Length |
|---|---|---|---|---|
| 5' Ala1 | $tRNA^{Ala}UGC$ | Northern blot | CTCTCCCAGCTGAGCTATAGCCCC | 24 mer |
| 3' Ala1 | $tRNA^{Ala}UGC$ | Northern blot | GGAGCTATGCGGGATCGAACCGC | 23 mer |
| 5' Arg2 | $tRNA^{Arg}ACG$ | Northern blot | CTATCCAGCTGAGCTACGGATGC | 23 mer |
| 3' Arg2 | $tRNA^{Arg}ACG$ | Northern blot | GCATCCGGGAGGATTCGAACCTC | 23 mer |
| 5' Leu1 | $tRNA^{Leu}CAG$ | Northern blot | GTCTACCAATTCCGCCACCTTCGC | 24 mer |
| 3' Leu1 | $tRNA^{Leu}CAG$ | Northern blot | CGAGGGGGGGACTTGAACCC | 21 mer |
| 5' Arg4 | $tRNA^{Arg}UCU$ | Northern blot | CGTTGCTCTATCCAACTGAGCTAAG | 25 mer |
| 3' Arg4 | $tRNA^{Arg}UCU$ | Northern blot | CGCGCCCTGCAGGATTCGAACCTGC | 25 mer |
| 34-mer | | Size marker | AUGUACACGGAGUCGAGCUCAACCCGCAACGCGA (Phos) | 34 mer |
| 54-mer | | Size marker | GGTCATCCGGCGACAAAAACAAAGTTGTCGGTTTTGTGTTTAGGGATCCCCGGG | 54 mer |

**Suppl. Table 1. Oligonucleotide sequences used in this study.** Deoxy oligonucleotides were [$^{32}$P]-labeled at the 5’-end and used in Northern blot assays to reveal tsRNAs (Varshney et al., 1991). tRNAs are identified by amino acid specificity, isoacceptor number and anticodón sequence.

**Varshney, U., Lee, C. P. and RajBhandary, U. L.** Direct analysis of aminoacylation levels of tRNAs in vivo. Application to studying recognition of Escherichia coli initiator tRNA mutants by glutaminyl-tRNA synthetase. *J Biol Chem*. **(1991)**266:24712-24718

| tRNA | tRFs and tRNA halves (tsRNAs) from ribosome-free | tRFs and tRNA halves (tsRNAs) from 30S | tRNA 5´halves from ribosome-free | tRNA 5'halves from 30S | tRNA 3´halves from ribosome-free | tRNA 3'halves from 30S |
|---|---|---|---|---|---|---|
| Ala1(UGC) | 34879 | 7099 | 18089 | 1288 | 15373 | 3937 |
| Ala2(GGC) | 33898 | 4147 | 24360 | 1929 | 7730 | 1309 |
| Arg2(ACG) | 13030 | 2197 | 7031 | 424 | 4674 | 693 |
| Arg3(CCG) | 1657 | 578 | 1127 | 68 | 285 | 59 |
| Arg4(UCU) | 395 | 211 | 224 | 21 | 86 | 57 |
| Arg5(CCU) | 1700 | 203 | 182 | 5 | 1467 | 176 |
| Asn (GUU) | 323409 | 14228 | 307301 | 5620 | 811 | 635 |
| Asp1(GUC) | 57137 | 8726 | 21913 | 1792 | 30665 | 2755 |
| Cys (GCA) | 26328 | 2598 | 22533 | 1606 | 2233 | 182 |
| Gln1(CUG) | 24900 | 2272 | 1623 | 145 | 9297 | 2657 |
| Gln2(UUG) | 11237 | 3286 | 1251 | 319 | 22932 | 1444 |
| Glu (UUC) | 3811 | 3743 | 1553 | 709 | 1781 | 1858 |
| Gly1 (CCC) | 30754 | 2753 | 16250 | 746 | 8831 | 501 |
| Gly2 (UCC) | 4766 | 891 | 2006 | 146 | 2438 | 246 |
| Gly3 (GCC) | 22458 | 3775 | 4274 | 277 | 16220 | 2034 |
| His (GUG) | 39531 | 5598 | 7117 | 3133 | 31515 | 1832 |
| Ile1(GAU) | 15208 | 2390 | 8988 | 591 | 555 | 56 |
| Ile2(CAC) | 4 | 3 | 1 | 0 | 85 | 0 |
| Ile2(CAU) | 37 | 61 | 2 | 0 | 85 | 2 |
| Leu1(CAG) | 18783 | 12532 | 10421 | 1596 | 4781 | 3336 |
| Leu2(GAG) | 3713 | 1955 | 860 | 612 | 424 | 385 |
| Leu3(UAG) | 1960 | 416 | 300 | 71 | 424 | 82 |
| Leu4(CAA) | 0 | 0 | 0 | 0 | 0 | 0 |
| Leu5(UAA) | 1656 | 804 | 313 | 105 | 424 | 324 |
| Lys (UUU) | 1067 | 936 | 222 | 39 | 334 | 58 |
| Met f1(CAU) | 6395 | 1783 | 2691 | 369 | 2863 | 244 |
| Met f2(CAU) | 1153 | 369 | 632 | 165 | 424 | 76 |
| Met m (CAU) | 1144 | 615 | 536 | 177 | 29 | 3 |
| Phe (GAA) | 522 | 408 | 33 | 24 | 4 | 2 |
| Pro1(CGG) | 11683 | 2144 | 9365 | 1488 | 1306 | 174 |
| Pro2(GGG) | 8344 | 640 | 7103 | 364 | 1050 | 103 |
| Pro3(UGG) | 3295 | 1496 | 1090 | 146 | 1532 | 699 |
| SelC(UCA) | 4600 | 428 | 2410 | 202 | 2095 | 158 |
| Ser1(UGA) | 582 | 746 | 149 | 23 | 230 | 52 |
| Ser2(CGA) | 2453 | 380 | 1148 | 121 | 1206 | 109 |
| Ser3(GCU) | 5771 | 3503 | 4886 | 500 | 627 | 303 |
| Ser5(GGA) | 15950 | 4148 | 12003 | 2272 | 3676 | 397 |
| Thr2(CGU) | 8770 | 687 | 144 | 109 | 8391 | 457 |
| Thr1y3(GGU) | 13449 | 1607 | 11173 | 474 | 1736 | 412 |
| Thr4(UGU) | 11135 | 5010 | 6026 | 1477 | 3795 | 1514 |
| Trp (CCA) | 1544 | 1026 | 447 | 80 | 193 | 50 |
| Tyr (GUA) | 2583 | 1721 | 1284 | 283 | 686 | 441 |
| Val1(UAC) | 728 | 1938 | 75 | 532 | 403 | 315 |
| Val2(GAC) | 26353 | 1825 | 20131 | 989 | 5269 | 424 |
| **Total** | **798772** | **111876** | **539267** | **31037** | **198965** | **30551** |

**Suppl. Table 2. Reads for tRNA halves and tRFs in the 30S and ribosome-free fractions.**The *Escherichia coli* tRNAs, identified by sequence homology, are listed in the first column. Columns 2 and 3 list the total reads for tsRNAs for each tRNA (used to elaborate Fig.4C). Columns 4 to 7 list the reads for 5' and 3' tRNA halves from the indicated 30S or ribosome-free fractions (used to elaborate Suppl. Fig. 3C and D). All data are from the long-read RNAseq experiment. tRNAs are identified by amino acid specificity, isoacceptor number and anticodón sequence.

| | Ribosome-free<br>30S 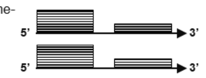 | Ribosome-free<br>30S 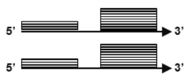 | Ribosome-free<br>30S 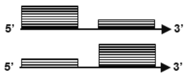 | Ribosome-free<br>30S 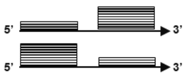 |
|---|---|---|---|---|
| 1 | Ala2(GGC)$^{alaW}$ | Arg5(CCU)$^{argW}$ | Ala1(UGC)$^{alaT}$ | His(GUG)$^{hisR}$ |
| 2 | Ala2(GGC)$^{alaX}$ | Asp1(GUC)$^{aspT}$ | Ala1(UGC)$^{alaU}$ | Leu2(GAG)$^{leuU}$① |
| 3 | Asn(GUU)$^{asnT}$ | Asp1(GUC)$^{aspU}$ | Ala1(UGC)$^{alaV}$ | Met f1(CAU)$^{metV}$① |
| 4 | Asn(GUU)$^{asnU}$ | Asp1(GUC)$^{aspV}$ | Arg2(ACG)$^{argQ}$ | Met f1(CAU)$^{metW}$① |
| 5 | Asn(GUU)$^{asnV}$ | Gln2(UUG)$^{glnU}$ | Arg2(ACG)$^{argV}$ | Met f1(CAU)$^{metZ}$① |
| 6 | Asn(GUU)$^{asnW}$ | Gln1(CUG)$^{glnV}$① | Arg2(ACG)$^{argY}$ | Ser2(CGA)$^{serU}$ |
| 7 | Cys(GCA)$^{cysT}$ | Gln2(UUG)$^{glnW}$ | Arg2(ACG)$^{argZ}$ | Val1(UAC)$^{valU}$ |
| 8 | Gly1(CCC)$^{glyU}$ | Gln1(CUG)$^{glnX}$ | Arg3(CCG)$^{argX}$① | Val1(UAC)$^{valX}$ |
| 9 | Ile1(GAU)$^{ileT}$ | Glu2(UUC)$^{gltT}$①② | Arg4(UCU)$^{argU}$① | val1(UAC)$^{valY}$ |
| 10 | Ile1(GAU)$^{ileU}$ | Glu2(UUC)$^{gltU}$①② | Leu1(CAG)$^{leuT}$ | Val1(UAC)$^{valT}$① |
| 11 | Ile1(GAU)$^{ileV}$ | Glu2(UUC)$^{gltV}$①② | Thr1,3(GGU)$^{thrT}$① | val1(UAC)$^{valZ}$①② |
| 12 | Metm(CAU)$^{metT}$ | Glu2(UUC)$^{gltW}$①② | Thr4(UGU)$^{thrU}$ | |
| 13 | Metm(CAU)$^{metU}$ | Gly2(UCC)$^{glyT}$ | | |
| 14 | Met f2(CAU)$^{metY}$①② | Gly3(GCC)$^{glyV}$① | | |
| 15 | Pro1(CGG)$^{proK}$ | Gly3(GCC)$^{glyW}$ | | |
| 16 | Pro2(GGG)$^{proL}$ | Gly3(GCC)$^{glyX}$① | | |
| 17 | Sec(UCA)$^{selC}$ | Gly3(GCC)$^{glyY}$① | | |
| 18 | Ser3(GCU)$^{serV}$①② | Leu1(CAG)$^{leuP}$ | | |
| 19 | Ser5(GGA)$^{serW}$ | Leu1(CAG)$^{leuQ}$ | | |
| 20 | Ser5(GGA)$^{serX}$ | Leu1(CAG)$^{leuV}$ | | |
| 21 | Thr1,3(GGU)$^{thrV}$ | Leu3(UAG)$^{leuW}$ | | |
| 22 | Trp(CCA)$^{trpT}$①② | Leu5(UAA)$^{leuZ}$ | | |
| 23 | Tyr(GUA)$^{tyrU}$ | Lys(UUU)$^{lysQ}$ | | |
| 24 | Val2(GAC)$^{valV}$ | Lys(UUU)$^{lysT}$ | | |
| 25 | Val2(GAC)$^{valW}$ | Lys(UUU)$^{lysW}$ | | |
| 26 | | Lys(UUU)$^{lysY}$ | | |
| 27 | | Lys(UUU)$^{lysV}$ | | |
| 28 | | Lys(UUU)$^{lysZ}$ | | |
| 29 | | Pro3 (UGG)$^{proM}$ | | |
| 30 | | Ser1(UGA)$^{serT}$ | | |
| 31 | | Thr2(CGU)$^{thrW}$ | | |
| 32 | | Tyr(GUA)$^{tyrT}$ | | |
| 33 | | Tyr(GUA)$^{tyrV}$ | | |

**Suppl. Table 3. Relative abundance of 5’ and 3’ reads of halves for 30S and ribosome-free fractions.** Based on G-browser images of RNAseq, the profiles were separated in four groups according with the relative abundance of 5’ and 3’ end reads. The arrows at the top of each column indicate the orientation 5’ to 3’ of the tRNA genes. Line stacks above the arrows represent the relative abundance of reads for 5’ and 3’ halves. The gap between the stacks indicates processing probably at the anticodon loop. ①: tsRNAs including a central fragment in the ribosome free fraction and ②: tsRNAs including a central fragment in the 30S fraction. Genes encoding tRNAs Ile2, Leu4, and Phe are not listed because they show scarce or no reads (see Suppl. Table 1). leu4 gene may be absent by a deletion in strain CP78 (see text). The tRNAs are identified by amino acid specificity., anticodon number and gene letter.

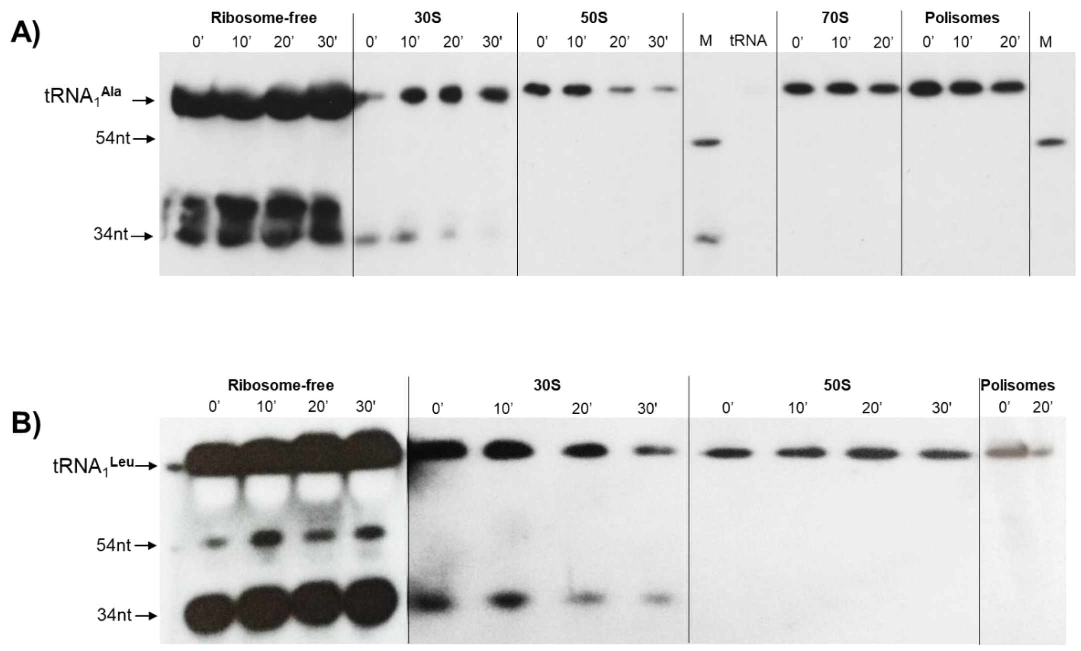


**Suppl. Fig. 4. Arrest of protein synthesis of a thermosensitive Pth mutant gradually reduces accumulation of tRNA fragments.** *E. coli* AA7852, a Pth$^{ts}$ variant of CP78, was grown at 32°C to log phase, the culture was shifted to non-permissive temperature (43°C), and samples were collected at the indicated times. For each time sample the crude extracts were ran through sucrose gradients and the fractions collected, resolved though acid urea PAGE and Northern blotted with [$^{32}$P] 5´-tRNA$^{Ala}$UGC and **B)** with [$^{32}$P] 5´-tRNA$^{Leu}$CAG. Size markers were deoxi-oligonucleotides of the indicated size.
